# A two-step selection method for in vitro evolution of translational proteins

**DOI:** 10.64898/2026.05.09.724044

**Authors:** Akari Sakurai, Kentaro Shoji, Norikazu Ichihashi

**Affiliations:** Department of Life Science, Graduate School of Arts and Sciences, The University of Tokyo, 3-8-1 Komaba, Meguro-ku, Tokyo 153-8902, Japan; Komaba Institute for Science, The University of Tokyo, 3-8-1 Komaba, Meguro, Tokyo 153-8902, Japan; Research Center for Complex Systems Biology, Universal Biology Institute, The University of Tokyo, 3-8-1 Komaba, Meguro, Tokyo 153-8902, Japan

**Keywords:** PURE system, reconstituted translation system, in vitro evolution, directed evolution, EF-G

## Abstract

Improving the reconstituted translation system is a key requirement for bottom-up synthetic biology. Here, we developed a two-step in vitro evolutionary method that can be used for improving translational proteins. In this method, two distinct conditions were sequentially applied while maintaining genotype-phenotype linkage in water-in-oil droplets. Using this method, we performed in vitro evolution of four translation factors, IleRS, PheRS, EF-G, and EF-Tu, and identified mutations that modestly enhanced translation activity in in vitro expression assays. One of the EF-G mutations (P610S) increased activity per protein approximately 2-fold for the recombinant protein purified from E. coli. This selection method is useful for improving translational proteins for bottom-up synthetic biology.

## Introduction

The reconstituted translation system, commonly referred to as the PURE system, is a powerful platform for bottom-up synthetic biology. Numerous genes have been expressed in the PURE system to construct various biological phenomena in vitro, including genetic circuits, artificial cells, and cell-free protein synthesis and engineering, as extensively reviewed elsewhere^1–6^. However, the translational yield of the PURE system remains significantly lower than that of cell extract-based systems^7^, thereby limiting its applications. Improving translation efficiency in the PURE system is the next critical challenge.

Several strategies have been explored to enhance translation activity, including the addition of auxiliary proteins^7–12^, optimization of system composition^13–15^, control of redox conditions^16^, replacement with heterologous translational components^17^, and the use of dialysis systems^15,18–21^. However, in vitro evolution of translational proteins has not yet been explored, probably due to the lack of methodology. Because currently used proteins are optimized for intracellular conditions through the long history of evolution of life, in vitro evolution may yield variants better suited to cell-free environments.

A major obstacle to the in vitro evolution of translation factors (TFs) is that expression of a target TF during selection requires supplementation with the same TF, creating background activity that interferes with selection. Since selection experiments typically involve low gene copy numbers within compartments, this background can severely reduce selection efficiency. Indeed, as shown in this study, conventional one-step selection failed to achieve efficient enrichment.

To address this issue, we developed a two-step selection method in water-in-oil droplets, based on previous transcription/translation-coupled DNA replication system in vitro^22^. Using this approach, we performed selection experiments on four TFs (IleRS, PheRS, EF-G, and EF-Tu) and identified some mutations that enhance translation activity. The two-step selection method provides a useful strategy for improving the reconstituted translation systems.

## Results and Discussion

The workflow of our new two-step selection strategy is shown in Fig. 1A. In the first reaction, droplets contain circular DNA encoding a target TF and a PURE system supplemented with a reduced concentration of the target TF. The target protein is expressed under conditions optimized for translation. The droplets are then diluted tenfold with a second set of droplets containing linear DNA encoding phi29 DNA polymerase (DNAP), dNTPs, and a PURE system lacking the target TF and optimized for DNA replication. Droplet fusion is induced by gentle mixing. In the second reaction, DNAP is expressed using the TF produced in the first step and replicates the circular DNA via rolling-circle replication. The replicated DNA is then recovered, amplified by PCR, circularized, and used in the next round. A key advantage of this method is that buffer conditions can be changed between the 1^st^ and 2^nd^ reactions to a certain extent, and thus, the 1^st^ and 2^nd^ reactions were optimized for translation and DNA replication, which require substantially different conditions^23–26^, respectively in this study.

**Figure 1.**
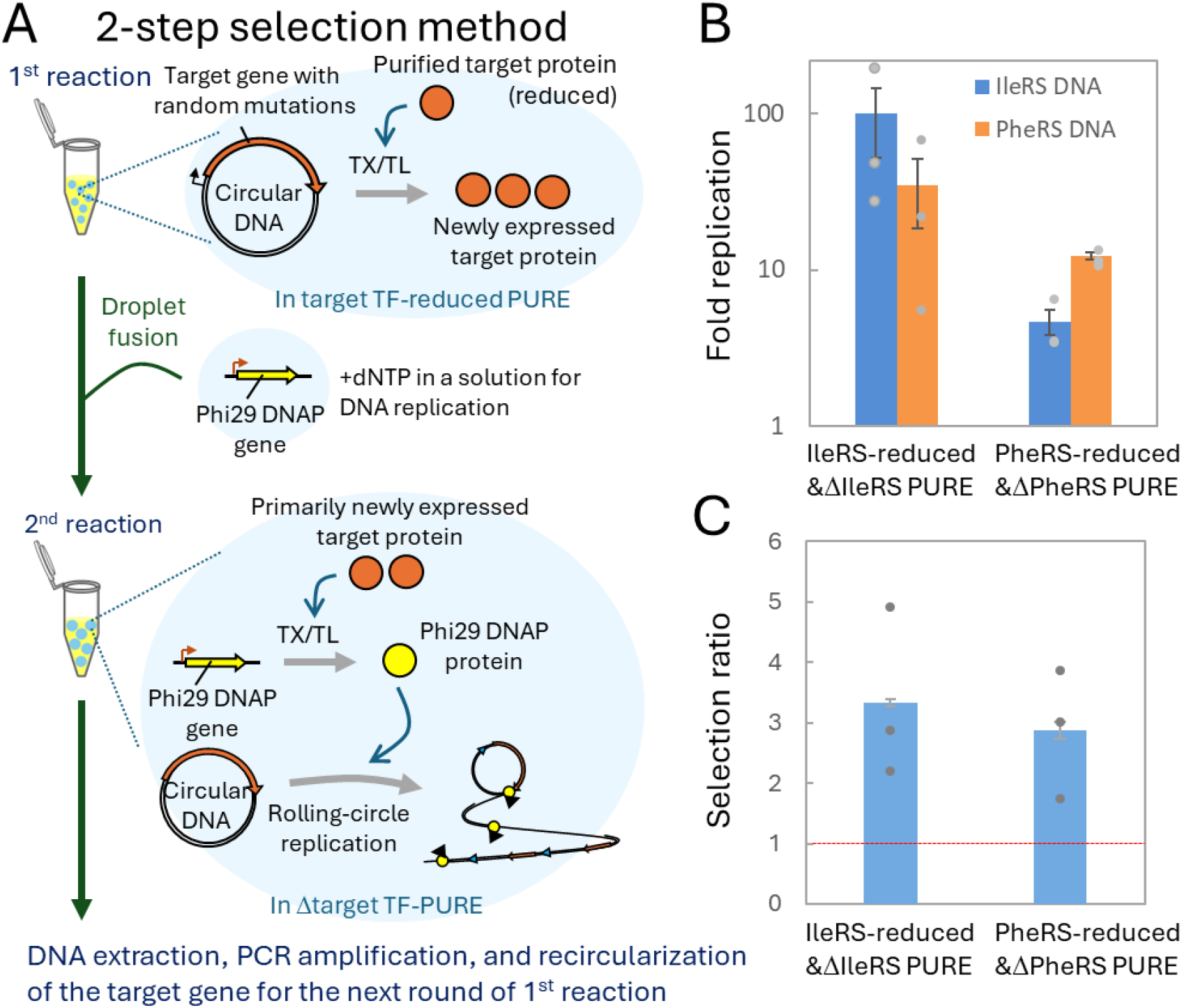
Two-step selection method for TF evolution. A) Schematic of the two-step selection method. All reaction mixtures were encapsulated in water-in-oil droplets. In the 1^st^ reaction, the target TF protein was expressed from a circular DNA in a PURE system containing a reduced concentration of the target TF. Then, an aliquot of droplets was fused with a 10-fold volume of fresh droplets that contained linear DNA encoding the phi29 DNAP gene and dNTPs. In the second reaction, DNAP was expressed using the target TF produced in the 1^st^ reaction and catalyzed rolling-circle replication of the circular DNA encoding the target TF. The replicated DNA was extracted from the droplets, amplified by PCR, circularized, and used for the first reaction of the next round. B) Selection test for target TFs. A single round of the two-step selection was conducted with a mixture of two circular DNAs encoding PheRS or IleRS at equal molar ratios, under two different conditions for selecting IleRS (IleRS-reduced & ΔIleRS PURE) or PheRS (PheRS-reduced & ΔPheRS PURE). C) Selection efficiency. A red dotted line indicates the no-selection level. Average values from three independent experiments are shown with standard errors.

To evaluate selection efficiency, we performed a mock selection experiment using a mixture of IleRS- or PheRS-encoding DNAs at equal ratios. When we used IleRS-reduced PURE in the 1^st^ reaction and ΔIleRS PURE in the 2^nd^ reaction, aiming at IleRS selection, IleRS DNA replicated more than PheRS DNA (Fig. 1B). In contrast, when we used PheRS-reduced PURE in the 1^st^ reaction and ΔPheRS PURE in the 2^nd^ reaction, aiming at PheRS selection, PheRS DNA replicated more than IleRS DNA. The selection ratios (replication fold of targeted DNA relative to non-targeted DNA) were around 3 (Fig. 1C), indicating 3-fold enrichment of target DNA. In contrast, a conventional one-step method, in which expression and replication occur simultaneously (Fig. S1A), showed no significant enrichment (Figs. S1B and S1C), demonstrating the advantage of the two-step approach.

We repeated the two-step selection method for 10 rounds using four TFs (IleRS, PheRS, EF-G, and EF-Tu) as targets. IleRS and PheRS were chosen because they exhibit relatively low activities when expressed in the PURE system^17,27^ and are of similar sizes. EF-G and EF-Tu were chosen because they are more abundant in the PURE system, and improving their activities is beneficial. The initial circular DNA libraries were prepared using a low-fidelity PCR polymerase (KOD FX, Toyobo) to introduce mutations randomly, followed by self-ligation. For IleRS, PheRS, and EF-Tu, we performed duplicate experiments with a single reduced TF concentration in the 1^st^ reaction (3.6 nM IleRS, 1.2 nM PheRS, and 5 μM EF-Tu). For EF-G, we performed two experiments with two different EF-G concentrations in the 1^st^ reaction (30 or 100 nM) because DNA replication was low at the initially tested concentration (30 nM). These concentrations in the 1^st^ reaction were set such that protein concentration was linearly proportional to translation output (see Methods).

The replication fold at each round is shown in Fig. S2. The DNA for IleRS, PheRS, and EF-G replicated in most rounds, whereas EF-Tu DNA did not show clear replication across all rounds, implying that EF-Tu expression in the 1^st^ reaction was insufficient to support DNA replication in the 2^nd^ reaction. Although the low replication efficiency was discouraging, we continued the selection rounds, expecting the emergence of more active mutants.

After 10 rounds of selection, the DNA was subjected to deep sequencing using a short-read sequencer (DNBSEQ), and mutation frequencies were analyzed. The four most frequent mutations were extracted from each experiment as candidates for beneficial mutations (Table S1). From these mutations, we selected three to four nonsynonymous mutations for further analysis. Each of the selected mutations was introduced into the corresponding target TF gene, and their activities were assayed using the method shown in Fig. 2A. In this assay, each mutant TF was expressed in the first reaction in a PURE system containing reduced levels of the target TF, and then the reaction mixture was diluted with a second PURE system lacking the target TF but containing GFP-encoding DNA to drive GFP expression. We measured GFP fluorescence, which is expected to reflect the translation activity of the TFs expressed in the first reaction.

**Figure 2.**
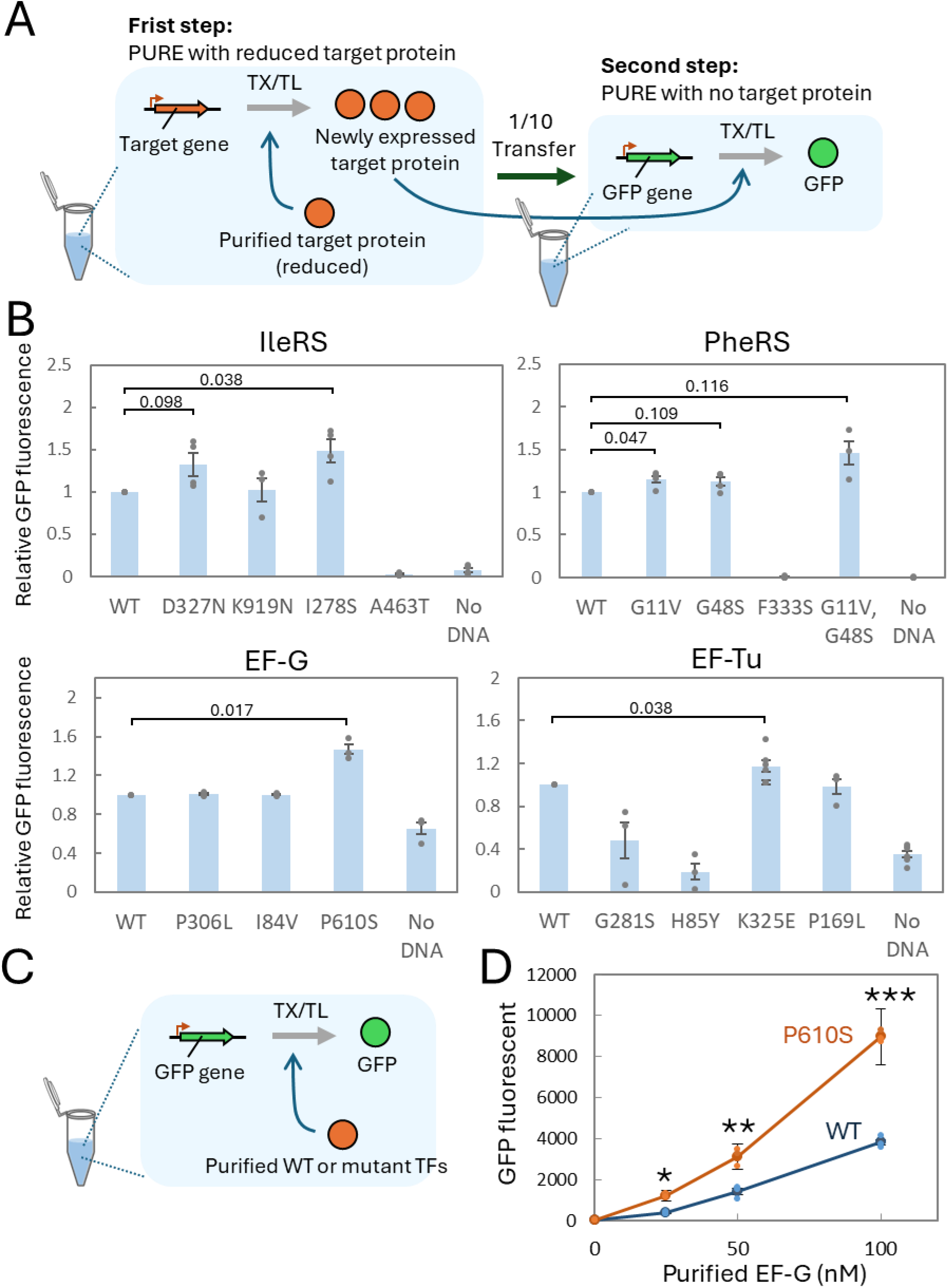
Effect of selected mutations on translation activity. (A) Schematic of the gene expression-coupled translation assay. In the first reaction, each mutated TF was expressed in a PURE system containing reduced levels of the corresponding TF, and then an aliquot (one-tenth volume) was transferred to a second PURE system lacking the target TF but containing a GFP-encoding gene. After incubation for 4 h at 30 °C, GFP fluorescence was measured. (B) GFP fluorescence relative to that of the wild-type TF. A reaction without the addition of the corresponding TF gene was used as a control (No DNA). Average values from 3–6 independent experiments are shown with standard errors. Statistical significance was determined using a one-sample t-test against a theoretical mean of 1.0, and the P values are indicated in the panel. C) Schematic of the translation assay using purified TFs. GFP expression was measured by fluorescence in a PURE system containing various concentrations of wild-type or mutant TFs. D) GFP expression in the presence of wild-type or P610S mutant EF-G. Average values from three technical replicates are shown with standard errors. P values from a Student’s t-test are indicated as *P = 0.0011, **P = 0.0020, and ***P = 0.00026.

For IleRS, PheRS, and EF-G, some mutations showed slightly (1.1 – 1.5-fold) higher activities than the wild type (D327N and I278S for IleRS, G11V and G48S for PheRS, P610S for EF-G, and K325E for EF-Tu) (Fig. 2B). In addition, the combination of G11V and G48S in PheRS exhibited much higher activity. These results indicated that these mutations enhance the activity of each TF when expressed in the PURE system. However, it should be noted that, in this assay, the observed increase in activity may result from multiple factors, such as enhanced transcription, translation, or activity per protein.

To find mutations that enhance activity per protein and more useful, we purified mutant TFs that exhibited more than 1.4-fold higher activity than the wild type (I278S for IleRS, G11V and G48S for PheRS, and P610S for EF-G) as recombinant proteins expressed in E. coli (Fig. S3). The translation activities were measured in the corresponding PURE system lacking each target TF, using GFP as a reporter protein (Fig. 2C). The IleRS and PheRS mutants exhibited translation activities similar to those of the wild type (Fig. S4), whereas the EF-G mutant P610S exhibited approximately 2-fold higher activity at 25 – 100 nM (Fig. 2D). These results indicate that the IleRS and PheRS mutations were effective only when the proteins were expressed in the PURE system, whereas the P610S mutation enhances EF-G activity per protein when expressed in E. coli.

The P610S mutation is located at the terminus of domain IV, a region that inserts into the ribosomal decoding center. There is limited knowledge regarding the function of this site; another mutation at this site (P610L) is known to enhance resistance to aminoglycoside antibiotics, while having only a minor effect on translation speed in the absence of the antibiotics^28^. The detailed mechanism of the P610S mutation remains unknown and requires further investigation.

## Conclusions

In this study, we developed a two-step selection method to improve the translation activity of TFs and identified several mutations that modestly enhanced the translation activity of IleRS, PheRS, EF-G, and EF-Tu when expressed in the PURE system. These mutations are useful for achieving a self-regenerating translation system, one of the major challenges in bottom-up synthetic biology, because enhanced translation activity reduces the required expression levels. Furthermore, one EF-G mutation (P610S) improved translation activity per protein even as a purified recombinant protein. This EF-G mutation allows the same level of translation with a smaller amount of protein and can therefore reduce the cost of EF-G preparation. The two-step selection method developed in this study paves the way for developing TFs optimized for in vitro conditions.

The two-step selection method still has room for improvement. First, the improvements in translation activity are modest for all identified mutations, and some mutations exhibit reduced activity (e.g., A463T in IleRS and F333S in PheRS). These limitations are likely due to insufficient selection efficiency in our method. There remains a background selection pathway in which even mutant TF genes that have lost translation activity can be replicated using pre-supplemented TFs, although this effect was partially reduced by employing the two-step method. Additional strategies to further reduce the influence of pre-supplemented TFs would enhance selection efficiency. Another possible improvement lies in the DNA replication scheme. The rolling-circle replication used here is simple but has uncertainties in its initiation mechanism, and some mutations may be selected based on DNA replication efficiency rather than translation activity. Adopting alternative replication schemes, such as the native phi29 virus replication system^25^, may improve the effectiveness of the selection.

## Materials and Methods

### DNA preparation

Circular DNA encoding the target TF gene was prepared as follows. First, target genes were PCR-amplified with a T7 promoter using primers 1 and 2–4, which contain a HindIII site, and the corresponding plasmids, which were used for expression of PURE system components^29^, as templates. The PCR fragments were then purified using a spin column (FastGene Gel / PCR Extraction Kit, Nippongene) and digested with HindIII, followed by self-ligation using T4 DNA ligase (Takara) according to the manufacturer’s instructions. A linear DNA fragment encoding phi29 DNA polymerase was prepared by PCR using primers 5 and 6 with a plasmid (pET-DNAPevo56) as a template, which encodes the DNA polymerase from clone 6 described in our previous study^30^. Each mutant TF was constructed by introducing mutations via whole-plasmid PCR using mutagenic primers and the corresponding plasmid, followed by self-ligation using the In-Fusion cloning method (Takara). The constructs were verified by Sanger sequencing. The linear DNA fragments encoding each TF gene used in Fig. 2B were prepared by PCR using primers 7 and 8 with the corresponding plasmids as templates, followed by purification using the spin column. The linear DNA fragment encoding GFP was prepared using the same method with the GFP-encoding plasmid pET-g5tag. All primer sequences are shown in Table S3.

### Reconstituted translation system

All components of the reconstituted translation system (PURE system) used in this study were purified and reconstituted in our laboratory as previously described^17^. The protein and ribosome compositions were the same as those in our previous study^17^, except for the target TFs, which were reduced or omitted as indicated in the figure legends.

### Selection experiments

Two-step selection was performed as follows. For the 1^st^ reaction, a circular DNA (0.1 nM) was mixed with the reconstituted translation system containing reduced concentrations of each TF (3.6 nM IleRS, 1.2 nM PheRS, 30 or 100 nM EF-G, and 5 μM EF-Tu), and the composition of the non-protein components (Table S2) was optimized for translation^15^. The mixture (10 μL) was vigorously emulsified using a homogenizer at 16 krpm for 1 min at 4 °C in buffer-saturated mineral oil (1 mL) prepared as described previously^31^ to generate a water-in-oil emulsion. After incubation at 30 °C for 4 h, an aliquot (50 μL) of the emulsion was collected from the middle of the tube and gently layered onto another emulsion for the 2^nd^ reaction (450 μL), which had been prepared in advance by emulsifying a mixture (10 μL) containing the translation system and a linear DNA fragment encoding phi29 DNAP (1 nM) under the same conditions described above. The non-protein composition of the reconstituted translation system in the emulsion for the 2^nd^ reaction was optimized for DNA replication^23^ (Table S2). The mixture of the emulsions was gently inverted 20 times in a 1.5 mL tube to induce droplet fusion and incubated at 30 °C for 16 h for the 2^nd^ reaction. Before and after this incubation, aliquots of droplets were diluted 100-fold with 1 mM EDTA (pH 8.0), and the concentration of the DNAP gene was measured by quantitative PCR using primers 9–12 and 13 (Table S3). After the 2^nd^ reaction, 200 μL of emulsion was centrifuged at 20 × g for 5 min at 23 °C to precipitate water droplets. The precipitate was resuspended in buffer TE (10 μL) and mixed with AMPure XP beads (18 μL). After incubation for 15 min at 23 °C, the beads were washed with 70% ethanol, and DNA was eluted with buffer TE (10 μL). Using the purified DNA as a template, the target TF gene was PCR-amplified using primers 14 and 15. When unexpected bands were observed, gel extraction and re-amplification by PCR were performed. The PCR product was digested with HindIII, purified using a spin column, self-ligated using T4 DNA ligase at 16 °C for 16 h, and purified again using a spin column. The resultant circular DNA was used for the 1^st^ reaction of the next round of the two-step selection. The reduced target TF concentrations in the first reaction were set such that protein concentration was linearly proportional to translation output. The relationship between each TF concentration and translation is shown in Figs. 2D, S4, and S5.

The one-step selection was performed as follows. The reaction mixture contained circular DNA (0.01 nM), a linear DNA fragment encoding phi29 DNAP (1 nM), and the reconstituted translation system with reduced TFs (3.6 nM IleRS and 1.2 nM PheRS) of the same composition as that used in the second reaction for the two-step method (i.e., optimized for DNA replication) shown above.

### Sequence analysis

After 10 rounds of the two-step selection process, PCR-amplified TF genes were analyzed by DNBSEQ. The sequence reads were aligned using BWA, and mutations were identified using SAMtools, along with their frequencies. Approximately 8,000 reads were analyzed across the entire regions of all four genes.

### TF protein purification

The wild-type and mutant IleRS, PheRS, and EF-G, used in Figs. 2D and S4, were purified using a histidine tag, as described previously for other translation proteins^17^, except that the gel-filtration chromatography step was omitted.

### Assay of mutant TFs

Gene expression–coupled translation assays in Figs. 2A and 2B were conducted in two steps. In the first step, the reaction mixture contained a linear DNA fragment (5 nM) encoding a target gene, reduced concentrations of each target TF (1.8 nM IleRS, 1.2 nM PheRS, 100 nM EF-G, and 5 μM EF-Tu), T7 RNA polymerase (0.1 μM), and a reconstituted translation system optimized for translation (Table S2, the same composition as that used in the 1^st^ reaction of the two-step selection method). After incubation for 4 h at 30 °C, the reaction mixture was diluted 10-fold with a second reaction mixture, which lacked each target TF and contained a linear DNA fragment encoding GFP (3 nM), T7 RNA polymerase (0.05 μM), and the same reconstituted translation system. The reaction mixture was incubated at 30 °C for 8 h, and GFP fluorescence was measured every 10 min. Fluorescence at 8 h was used for analysis.

Assays of purified TF proteins in Fig. 2D and S4 were conducted using the same reconstituted translation system optimized for translation, together with a linear DNA fragment encoding GFP (1 nM) and T7 RNA polymerase (0.05 μM). The reaction mixture was incubated at 30 °C for 8 h, and GFP fluorescence was measured every 10 min. Fluorescence at 8 h was used for analysis.

## Supporting Information

Supporting Information includes supplemental figures (Figures S1 – S5) and a table (Table S1 – S3).

## Author contribution

AS, KS, and NI planned and performed the experiments. NI wrote the manuscript.

## Acknowledgement

We thank Ms. Kayo Aoyama and Ayu Saito for their technical support.

