## Supplemental Figures and Tables for "A two-step selection method for in vitro evolution of translational proteins"

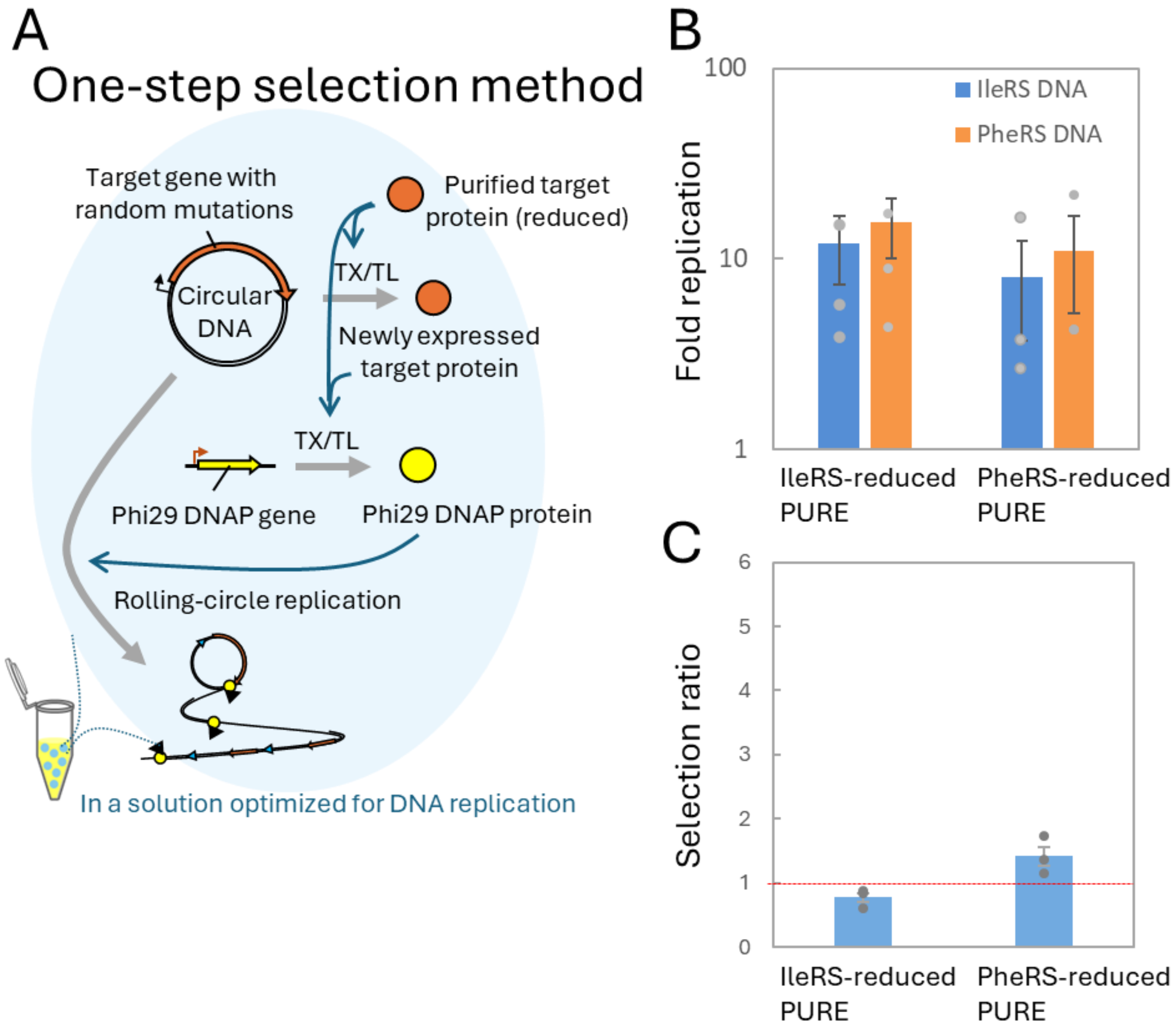


**Figure S1. One-step selection method.**

A) Schematic of the one-step selection method. All reaction mixtures were encapsulated in water-in-oil droplets. The reaction mixture contained circular DNA encoding a target TF, linear DNA encoding phi29 DNAP, dNTPs, and a PURE system containing a reduced concentration of the target TF (3.6 nM IleRS or 1.2 nM PheRS). In this reaction, the target TF protein was expressed from the circular DNA and was expected to catalyze DNAP expression from the linear DNA. The DNAP then catalyzed rolling-circle replication of the circular DNA encoding the target TF. B) Selection test for target TFs. A single round of the one-step selection was conducted with a mixture of two circular DNAs encoding PheRS or IleRS at equal molar ratios, under two different conditions for selecting IleRS (IleRS-reduced PURE) or PheRS (PheRS-reduced PURE). C) Selection efficiency. A red dotted line indicates the no-selection level. Average values from three independent experiments are shown with standard errors.

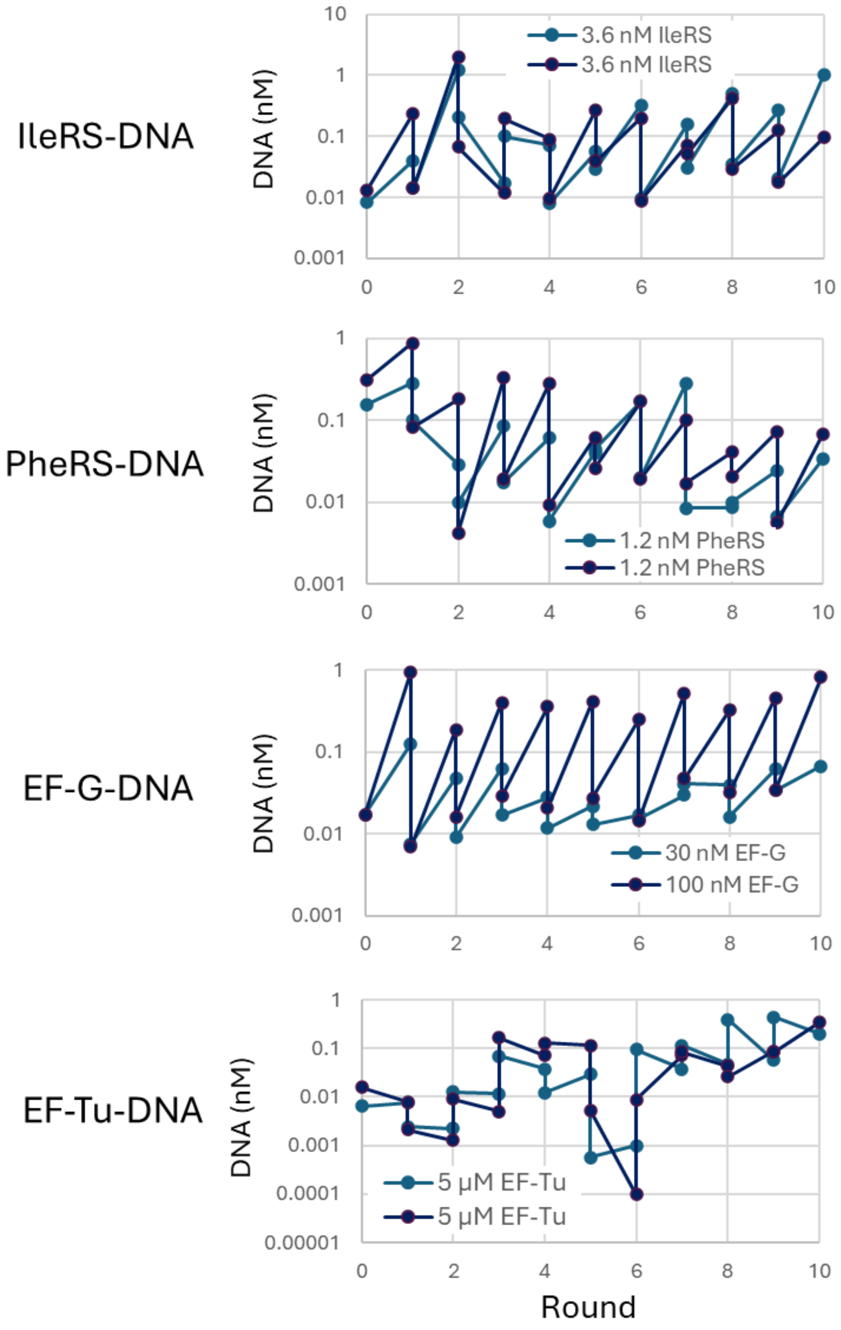


**Figure S2. Trajectories of replication fold at each round.**

The target gene concentrations were measured by quantitative PCR before and after the second reaction.

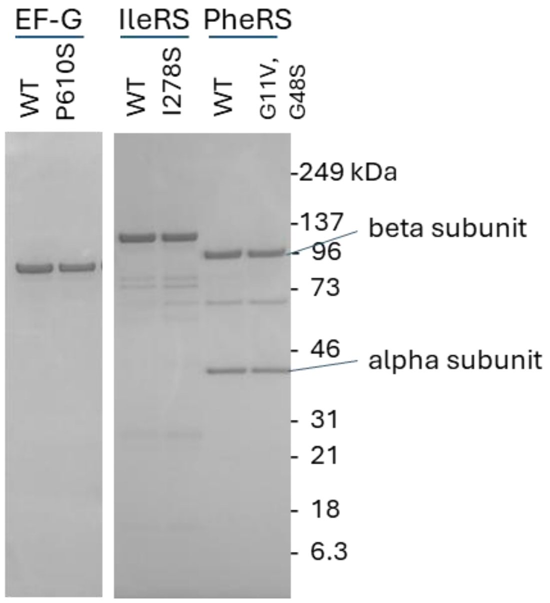


**Figure S3. SDS-PAGE analysis of purified proteins.**

Each purified protein (0.5 µg) was subjected to SDS-PAGE using a 10–20% gradient gel. Proteins were stained with Coomassie Brilliant Blue (CBB).

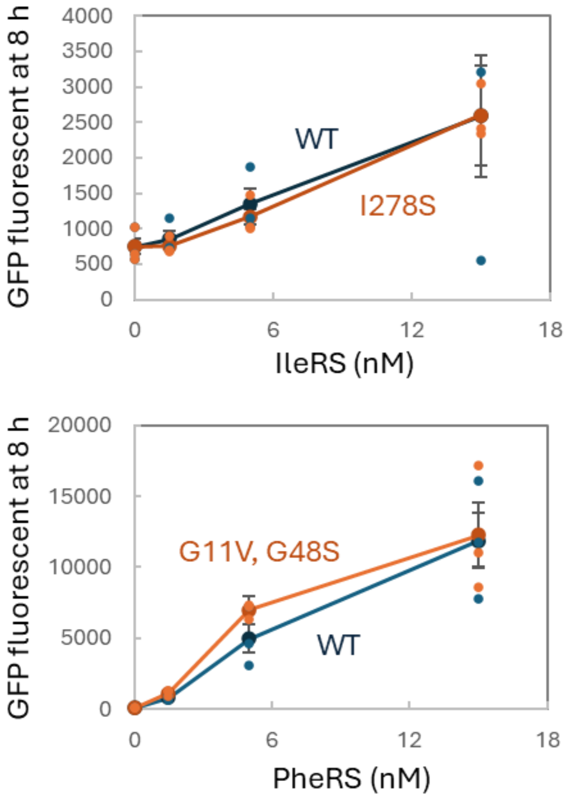


**Figure S4. GFP expression with purified TFs.**

The scheme of the translation assay is shown in Fig. 2C. GFP expression was measured by fluorescence in a PURE system containing various concentrations of wild-type or mutant TFs. Average values from three technical replicates are shown with standard errors.

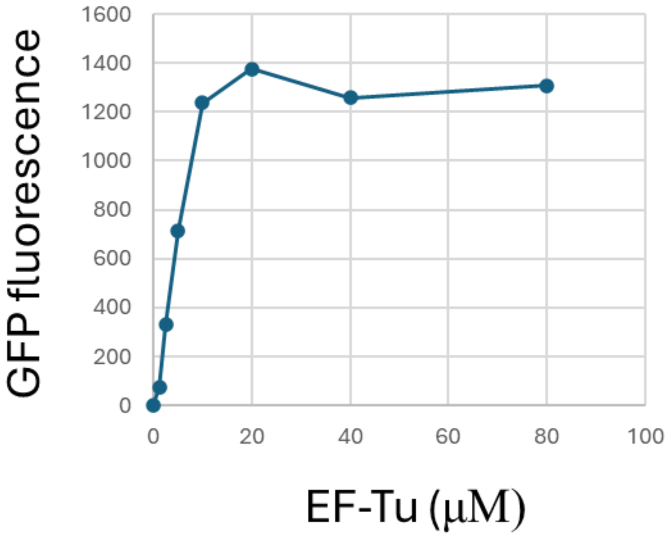


**Figure S5. GFP expression with purified EF-Tu.**

GFP expression was performed in the PURE system containing the indicated concentrations of wild-type EF-Tu at 37˚C for 2 h, and fluorescence was measured to determine the linear range.

**Table S1. Top four enriched mutations.** *Used for expression assay in Figure 2

| Target | Evolutionary condition | Mutation | Frequency |
|---|---|---|---|
| IleRS | 3.6 nM IleRS (#1) | G1086A (D327N)* | 0.377 |
| | | G2864T (K919N)* | 0.320 |
| | | T940G (I278S)* | 0.109 |
| | | G1494A (A463T)* | 0.088 |
| | 3.6 nM IleRS (#2) | G1086A (D327N)* | 0.320 |
| | | G2864T (K919N)* | 0.309 |
| | | G1494A (A463T)* | 0.086 |
| | | A2816G (E903E) | 0.085 |
| PheRS | 1.2 nM PheRS (#1) | G139T (G11V)* | 0.998 |
| | | G1301A (G48S)* | 0.143 |
| | | T1105C (F333S)* | 0.133 |
| | | G2198A (A347T) | 0.111 |
| | 1.2 nM PheRS (#2) | G139T (G11V)* | 0.997 |
| | | G1301A (G48S)* | 0.054 |
| | | G2198A (A347T) | 0.060 |
| | | C1372G (G71G) | 0.060 |
| EF-G | 30 nM EF-G | C1027T (P306L)* | 0.032 |
| | | G1065A (A319T) | 0.030 |
| | | T1748C (P546P) | 0.027 |
| | | G953A (A281A) | 0.027 |
| | 100 nM EF-G | A360G (I84V)* | 0.047 |
| | | C1938T (P610S)* | 0.040 |
| | | T946C (L279P) | 0.035 |
| | | G196A (R29H) | 0.034 |
| EF-Tu | 5 µM EF-Tu (#1) | C50T | 0.048 |
| | | G948A (G281S)* | 0.030 |
| | | A1080G (K325E)* | 0.027 |
| | | A1136G (E343E) | 0.026 |
| | 5 µM EF-Tu (#2) | T875C (C256C) | 0.030 |
| | | C613T (P169L)* | 0.028 |
| | | C360T (H85Y)* | 0.026 |
| | | G40A (G9C) | 0.023 |

**Table S2. Composition of non-protein components in the PURE systems**

| | Optimized for translation (used in 1st reaction in Fig. 1) | Optimized for DNA replication (used in 2nd reaction in Fig. 1) |
|---|---|---|
| Cys and Tyr | 0.3 mM each | 0.3 mM each |
| The other 18 amino acids | 0.36 mM each | 0.36 mM each |
| Hepes-KOH (pH 7.6) | 100 mM | 100 mM |
| Potassium glutamate | 280 mM | 70 mM |
| Spermidine | 1.5 mM | 0.375 mM |
| Magnesium acetate | 18 mM | 7.9 mM |
| Creatin phosphate | 25 mM | 25 mM |
| Dithiothreitol | 1.5 mM | 6 mM |
| 10-formyl-5.6.7.8.-tetrahydrofolic acid | 10 ng/µl | 10 ng/µl |
| ATP | 3.75 mM | 0.375 mM |
| GTP | 2.5 mM | 0.25 mM |
| CTP | 1.25 mM | 0.125 mM |
| UTP | 1.25 mM | 0.125 mM |
| EDTA (pH8.0) | 4 mM | - |
| tRNA (E. coli, Roche) | 3 µg/µl | 0.52 µg/µl |
| dNTP | - | 0.6 mM each |
| Pyrophosphatase, Inorganic (yeast, NEB) | - | 0.0002 unit/µl |
| T7 RNA polymerase | 0.05 – 0.1 µM | 0.0125 µM |

**Table S3. Primer list**

| | Sequence |
|---|---|
| Primer 1 | AAAATTAAGCTTGCGAAATTAATACGACTCACTATAGGG |
| Primer 2 for EF-G | tttataaagctTAAGCGAATGTTGCGAGCACGTCGACGGAGCTCGAATTCATTATTA |
| Primer 3 for IleRS | tttataaagctTAAGCGAATGTTGCGAGCACGCTTTCAGGCAAACTTACGTTTTTC |
| Primer 4 for PheRS and EF-Tu | TATATAAAGCTTaagcgaatgttgcgagcacGTCGACGGAGCTCGAATTC |
| Primer 5 | GCGAAATTAATACGACTCACTATAGGG |
| Primer 6 | GGTTATGCTAGTTATTGCTCAGCGG |
| Primer 7 | GCGTCCGGCGTAGAGGATC |
| Primer 8 | TCCGGATATAGTTCCTCCTTTCAG |
| Primer 9 for EF-G | CTGACCAAAGGTCGTGCATC |
| Primer 10 for PheRS | CCGTACACTCGAAGAAGAGGAG |
| Primer 11 for IleRS | GCACTACACCCAGGATGTC |
| Primer 12 for IleRS | GGTATCAAAGAGACTCAGAAGTC |
| Primer 13 | TAAGCGAATGTTGCGAGCAC |
| Primer 14 | AAAATTAAGCTTGCGAAATTAATACGACTCACTATAGGG |
| Primer 15 | tttataaagctTAAGCGAATGTTGCGAGCAC |
